# Molecular effects of mannanase-hydrolyzed coprameal to intestinal immunity in the Pacific white shrimp *(Litopenaeus vannamei)*

**DOI:** 10.1101/2021.02.16.431529

**Authors:** Wanilada Rungrassamee, Sopacha Arayamethakorn, Nitsara Karoonuthaisiri, Shih-Chu Chen, Eric Chang, Samu Chan, Yasuhiko Yoshida, Motohiro Maebuchi, Masahisa Ibuki

## Abstract

To mitigate disease outbreak, an alternative approach through enhancing shrimp immunity was explored. Mannan oligosaccharides (MOS) have been previously reported to enhance shrimp immune system. Here, coprameal samples were digested with mannanase to yield MOS, namely, mannanase-hydrolyzed coprameal (MCM) and feasibility of MCM as shrimp immunostimulant in grow-out ponds was determined. Pacific white shrimp (*Litopenaeus vannamei*) were fed with the commercial diet containing 1% MCM as the MCM-supplemented group and compared to the non-MCM control diet. There was no significant difference in survival rates between the MCM-supplemented and the control groups throughout the 4-month-period of the trial (*p* > 0.05). Gene expression analysis in shrimp intestines revealed that the transcript levels of antimicrobial peptides (*anti-lipopolysaccharide factor isoform 1* (*alf1*), *penaeidin (pen3a)* and *crustin (crus))* and lysozymes *(lyz)* were not significantly different in the MCM-supplemented group. Meanwhile, *C-type lectin* and *toll-like receptor* transcript levels, whose gene products play roles as pattern recognition proteins, were significantly higher in a group fed with MCM for 2- and 4-month periods than those of the control group (*p* < 0.05). The increased transcript levels of *C-type lectin* and *toll-like receptor* provide evidence for potential implementation of MCM as feed supplement to modulate shrimp immune system.

## Introduction

Aquaculture production play important role to provide food sources for increasing global population. The global demand for shrimp and prawn production is approximately 70% of crustaceans (FAO 2018). This results in the rapid growth of cultivated shrimp production, in particularly, an expanding of Pacific white shrimp farming throughout many Asian countries. However, the shrimp industry continues to suffer from production losses due to the higher disease outbreak incidents. To mitigate shrimp diseases, much attention has been paid to development of feed additives with potential to enhance growth and immune performance as a mean to prevent disease outbreaks in aquaculture.

In mammalians, prebiotics, which are non-digestible carbohydrates, are shown to selectively stimulate growth of beneficial bacteria in host digestive systems, and consequently, providing health benefits to their host animals (Al-Sheraji et al. 2013; Davani-Davari et al. 2019; Manning & Gibson 2004). Recently, prebiotics as immunostimulants have been applied in aquaculture as a promising approach for growth and immune enhancement (Hasan et al. 2019; Luna-González et al. 2012; Sang et al. 2010; Staykov et al. 2007). Among those, mannan oligosaccharides (MOS), derived from yeast cell wall *(Saccharomyces cerevisiae)*, have been shown to improve growth performance and enhance immune system of many aquatic animals (Dimitroglou et al. 2010; Salem et al. 2015; Zhang et al. 2012; Zhang et al. 2020). In crustaceans, applications of MOS as feed additives has been reported to significantly improve the survival rate of crayfish *(Cherax tenuimanus)* (Sang et al. 2010), and enhance growth rates in tropic spiny lobster *(Panulirus ornatus)* (Do Huu & Jones 2014) and black tiger shrimp *(Penaeus monodon)* (Sang et al. 2014; Sang & Fotedar 2010)and freshwater crayfish *(Astacus leptodactylus)* (Mazlum et al. **2011**). The dietary MOS from yeast can significantly increase intestinal microvilli length, growth and percent survival rates in the Pacific white shrimp (Gainza & Romero 2020; Zhang et al. 2012)

While MOS used in animal feed additives are mostly derived from yeast, MOS can also be obtained from other natural sources with high mannan sugar contents such as copra meal (Hossain et al. 1996; Kraikaew et al. 2020; Pangsri et al. 2015). Copra meal is a by-product from the coconut milk industry, and it is an alternative source for MOS in a form of β-1, 4 mannooligosaccharides (Ariandi et al. 2015). The potential applications of MOS from copra meal as immunostimulants have been explored in various animals such as broilers (Prayoonthien et al. 2017; Sundu et al. 2012) and the Pacific white shrimp (Chen et al. 2020; Li et al. 2018; Rungrassamee et al. 2014). Particularly, MOS from copra meal has been reported to improve immune levels and survival rates in the Pacific white shrimp juvenile under a pathogen exposure to *Vibrio harveyi* (Rungrassamee et al. 2014), providing a piece of evidence for MOS as a promising shrimp feed additive. Here, we further explored feasibility of MOS through mannanase-hydrolysis of copra meal (MCM) as feed additives in the grow-out pond systems and determined molecular effects on immune gene expression. These findings will lead to further development of MCM (MOS from copra meal) as effective prebiotics for shrimp to maintain their intestinal immunity in the grow-out phase.

## Materials and methods

### Preparation of mannanase-hydrolyzed coprameal (MCM)

The copra meals were hydrolyzed using mannanase enzyme to yield mannanase-hydrolyzed coprameal (MCM). Briefly, the defatted copra meals were incubated with 0.2-0.6% mannanase solution (15,000unit/g minimum, Shin-Nihon Chemical CO., Ltd., Aichi, Japan) at 40-70 **°**C for 18-30 h, and then, dried 2-6 h to obtain MCM product. MCM content analysis was carried out using following methods. Mannobiose was determined by high-performance anion-exchange chromatography coupled with a pulsed amperometric detection (HPAEC-PAD) system, and constituent sugars were analyzed by surfuric hydrolysation. Ashes were determined by a dry-ashing procedure at 55 ± 50°C for 2 h and moisture was determined by oven drying method at 105±1°C for 4 h. Fat was analyzed by a diethyl ether extraction method (AOAC Method 920.39). Protein content was determined with a Kjeldahl method (Lynch & Barbano 1999).

### Animal facility and feeding trials

A group of 16-day-old Pacific white shrimp postlarvae was cultivated in the grow-out ponds at the animal facility (Terminalia Garden Aquatic development, Taiwan) (Fig. 1). Shrimp were reared in brackish water with 0.07-0.15% salt concentrations and divided into two groups: a control feed with commercial feed pellets (a control diet) and a MCM-supplemented group (a test diet) (Table 1). The test diet was formulated to contain 1% of MCM in the commercial feed pellets. Shrimp were fed under these diets and intestine samples (N_pooled_ = 6 with 5 replicates) were aseptically collected at 2- and 4-month of the feeding trial period. All tissue samples were stored in RNA*later*^®^ RNA Stabilization Solution (Ambion, USA) to preserve tissue integrity for RNA extraction for gene expression analysis. Water quality in ponds was measured every other day for temperature, pH, and dissolved oxygen and weekly for ammonia–nitrogen, nitritenitrogen, and alkalinity levels and maintained at the standard ranges for rearing Pacific white shrimp (Rungrassamee et al. 2013).

**Figure 1.**
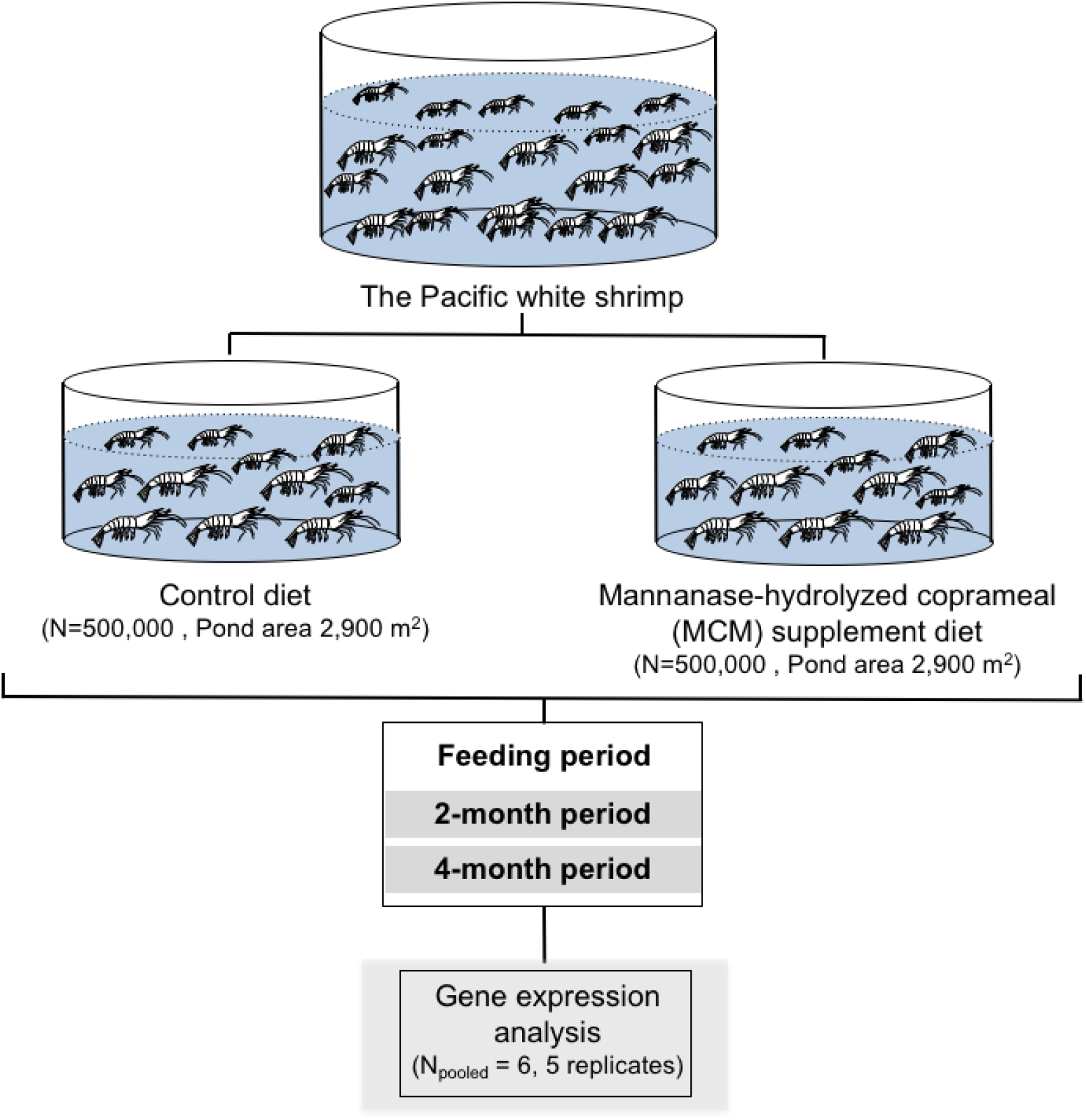
Experimental overview for evaluating effects of mannanase-hydrolysis of copra meal (MCM) as immunostimulant in Pacific white shrimp, *Litopenaeus vannamei* in a grow-out pond. The Pacific white shrimp postlarvae were divided into two groups: a control feed with commercial feed pellets and a 1% MCM-supplemented group. The intestine samples (N_pooled_ = 6 with 5 replicates) were collected at 2- and 4-month of the feeding trial period for gene expression analysis.

**Table 1.** Composition of feed diets used in this study.

| Raw material | % Feed composition |  |
| --- | --- | --- |
|  | Control diet | MCM supplement diet |
| Wheat flour | 35 | 35 |
| Fish meal | 20 | 20 |
| Squid meal | 10 | 10 |
| Water | 30 | 29 |
| Vitamin and yeast mix | 5 | 5 |
| MCM | 0 | 1 |

### Total RNA extraction and cDNA synthesis

Intestine tissues were homogenized in TRI REAGENT^®^ (Molecular Research Center, USA) for RNA extraction according to supplier’s instruction. RNA pellets were washed twice with 500 μL of 75% ice-cold ethanol, air-dried for 5 min, and dissolved in 50 μL diethylpyrocarbonate (DEPC)-treated water. To remove potential genomic DNA contamination, all RNA samples were treated with DNaseI enzyme (RQ1 RNase-free DNaseI, Promega, USA) for 60 min at 37 °C. The DNaseI treated-RNA samples were purified via phenol:chloroform extraction and precipitated with 1/10 volume of 3M sodium acetate and 1 volume of isopropanol as previously done (Rungrassamee et al. 2010). All treated-RNA samples were confirmed to be free from genomic DNA via PCR reaction. RNA concentration was quantified with a Nanodrop ND-8000 spectrophotometer (NanoDrop, UK). An ImProm-II™ Reverse Transcriptase System kit (Promega, USA) was used to synthesize the first strand cDNA using 1.5 μg of the DNA-free RNA sample as a template and each reaction was carried out according to the supplier’s protocol. The concentration of cDNA was measured by a Nanodrop ND-8000 spectrophotometer.

### Realtime PCR analysis of immune related genes transcript levels in response to MCM supplement diet

To determine MCM effects on expression levels of host immune genes, cDNA from each shrimp group after 2- and 4-month of the feeding trial periods were used as templates in realtime PCR analysis using a CFX96™ realtime system (Bio-rad Laboratories, USA). The transcript levels of immune-related genes chosen were antilipopolysaccharide factor1 *(alf1)*, crustin (*crus*), c-type lectin (c-*lec*), lysozyme (*lyz*), penaedin3 *(pen3a)* and Toll1 *(Toll1)* and the elongation factor-1 alpha *(EF1a)* was used as an internal control. Each realtime PCR reaction contained IQ^TM^ SYBR Green Supermix (Bio-Rad, USA), gene-specific primer pair (1.25 μM, Table 2) and cDNA (100 ng). The realtime PCR cycling parameters used were 95 °C for 3 min, 40 cycles of at 95 °C for 30 sec, 57 °C for 20 sec and 72 °C for 30 sec. The fluorescent signal intensities were recorded at the end of each cycle. Melting curve analysis was performed from 55°C to 95 °C with continuous fluorescence measurement at every 0.5 °C increment. Relative abundance of the target immune genes in intestines of the Pacific white shrimp was determined by the ΔΔct method (Livak & Schmittgen 2001). The relative fold change for each target gene was compared between the MCM-supplement to the control groups at the same time point.

**Table 2.**
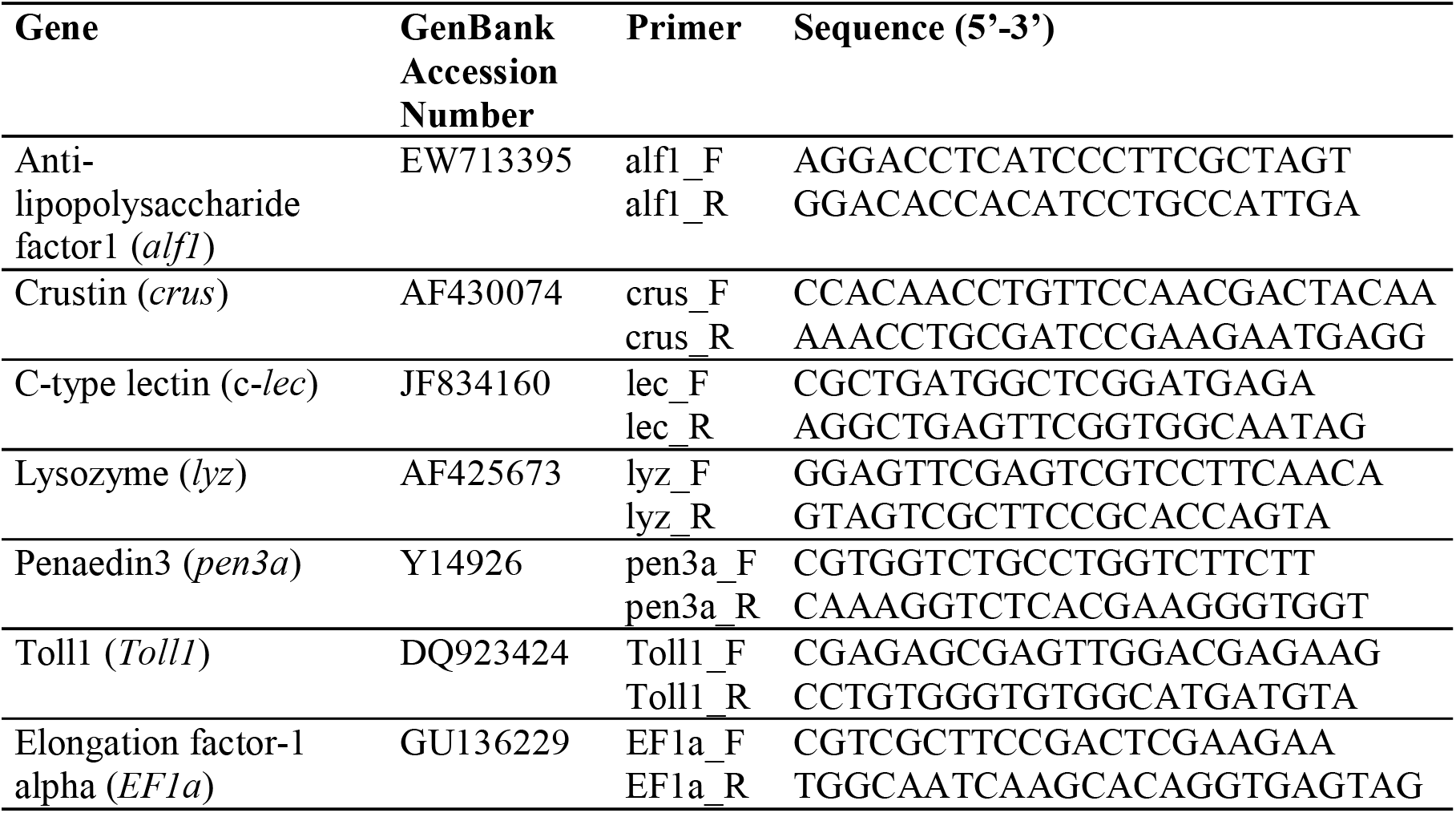
Primer sequences used in realtime PCR analysis

### Statistical analysis

Statistical analysis was conducted using SPSS of Windows version 15.0 to perform Student’s t test analysis for significant differences in shrimp weights or gene expression analysis (Landau & Everitt 2004).

## Results and discussion

### MCM component analysis

In this work, MOS supplement was derived in a form of mannanase-hydrolyzed coprameal (MCM). MCM yield was ~95% of the copra meal. The content analysis of MCM revealed high composition of carbohydrates (55.0%), followed by crude proteins (22.9%), fats (9.3%), moisture (7.0%) and ashes (5.8%) (Table 3). The free sugar contents in MCM were mainly mannobiose sugars (10.3%) and others were glucose, mannose, fructose, sucrose and mannotriose. Among those sugars, mannobiose has been reported for their important role as an immune modulator (Patel & Goyal 2012; Tiwari et al. 2020). For instance, mannobiose is able to activate innate immune response in chicken under *in vivo* and *in vitro* studies (Agunos et al. 2007; Duarte et al. 2014; Ibuki et al. 2010; Ibuki et al. 2011). This suggests that MCM supplementation in this study could potentially be used to enhance disease protection in animals.

**Table 3.**
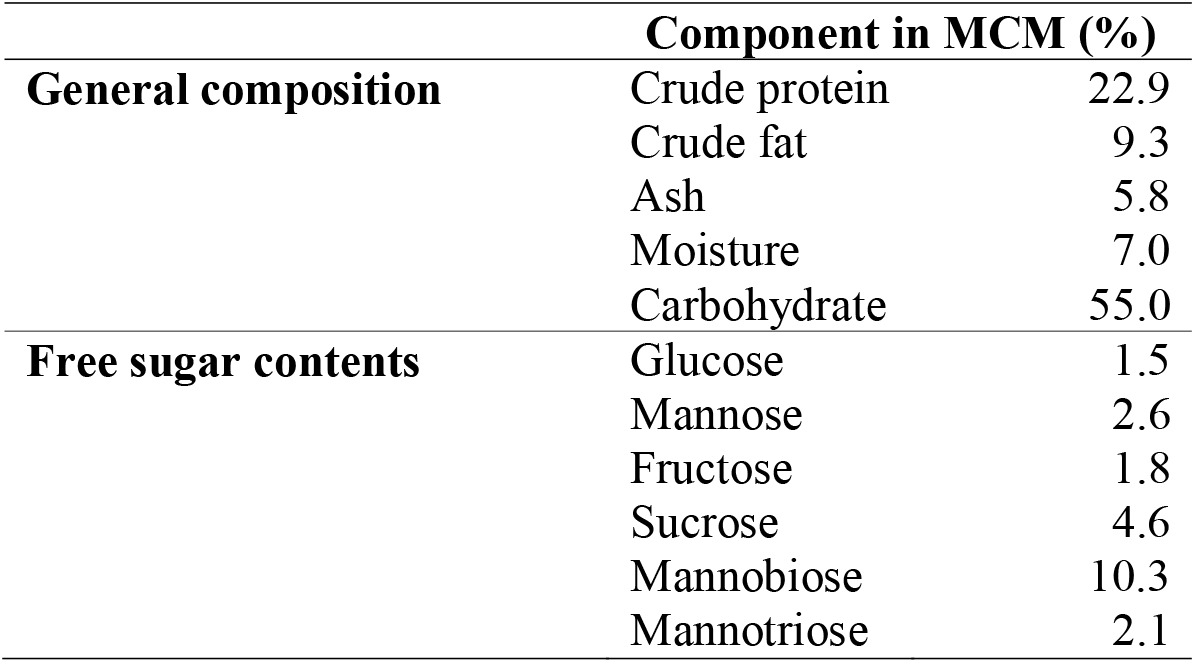
Content analysis of mannanase-hydrolyzed coprameal (MCM).

### Expression analysis of the immune related in response to MCM supplementary diet

Since shrimp digestive system, especially its intestine can be prone to pathogen invasion (Aguirre Guzman et al. 2010; Soonthornchai et al. 2015), it is important to determine approaches to modulate intestinal immune level. Hence, we selected genes encoding the antimicrobial peptides *(alf1, pen3a* and *crus)*, lysozyme (*lyz*), C-type lectin *(c-lectin)*, and Toll-like receptor protein *(Toll1)* as our genes of interest due to their important roles in digestive tract of the shrimp as part of host defense mechanisms against pathogens (Tassanakajon et al. 2013). To evaluate molecular effects of MCM supplement on host immune system, the relative expression levels of the aforementioned transcripts were compared in shrimp intestines at 2-month and 4-month after feeding with the MCM-supplement diet to those fed with the non-MCM as a control diet (Fig. 2). Our result showed that the MCM as dietary supplement did not affect the transcript levels of *alf1, pen3a* and *crus.* Additionally, the group fed with MCM showed a significant decreased in transcript level of *lyz* at the 2-month of the MCM-feeding period (p < 0.05), however the transcript level of *lyz* was not significantly different to the non-MCM diet at 4-month-period. This suggests that MCM did not have a direct effect on modulating antimicrobial peptides including lysozyme. In contrast, the *c-lectin* and *Toll1* expression levels were significantly higher than the control diet by ~2-fold when given MCM supplement diet at 2- and 4-month periods (p < 0.05). This suggests that MCM supplement specifically increases expression levels of *c-lectin* and *Toll1*, whose gene products play role as pattern recognition proteins. The encoded C-type lectin is a member of lectin family proteins that has been reported to bind to carbohydrate in a calcium-dependent manner (Weis et al. 1998). It plays important role in pathogen recognition by detecting conserved pathogen cell wall components of Gram-negative bacteria and further induces immune response to invaders (Bi et al. 2020; Medzhitov & Janeway 2002; Wang & Wang 2013). Lectins show strong bacterial-agglutination and opsonic activity, which facilitate phagocytosis (Luo et al. 2006). In the Chinese white shrimp, C-type lectin has been demonstrated *in vivo* to mediate in *V. anguillarum* clearance (Wang et al. 2009). Thus, the induction of *C-lectin* could provide a local immune response and protection in Pacific white shrimp intestine against bacterial pathogen invasion. Similarly, *Toll1*, encodes for Toll-like receptor protein, was also significantly increased at 2- and 4-month of the MCM supplemented group in comparison to the control group. Toll-like receptor proteins have been reported in mammals to play roles in viral pathogen detection and mediate responses to those viruses(Takeuchi & Akira 2009). Interestingly, the function of *Toll1* in the Pacific white shrimp has not yet been linked to viral diseases protection in shrimp (Labreuche et al. 2009), but it has been shown to involve in responses to bacterial pathogen (Liu et al. 2016). This suggests that MCM is a promising candidate to increase disease resistance to bacterial pathogens in Pacific white shrimp farming. Further investigation on optimal dosages and formulation for the MCM supplement should be conducted for future application.

**Figure 2.**
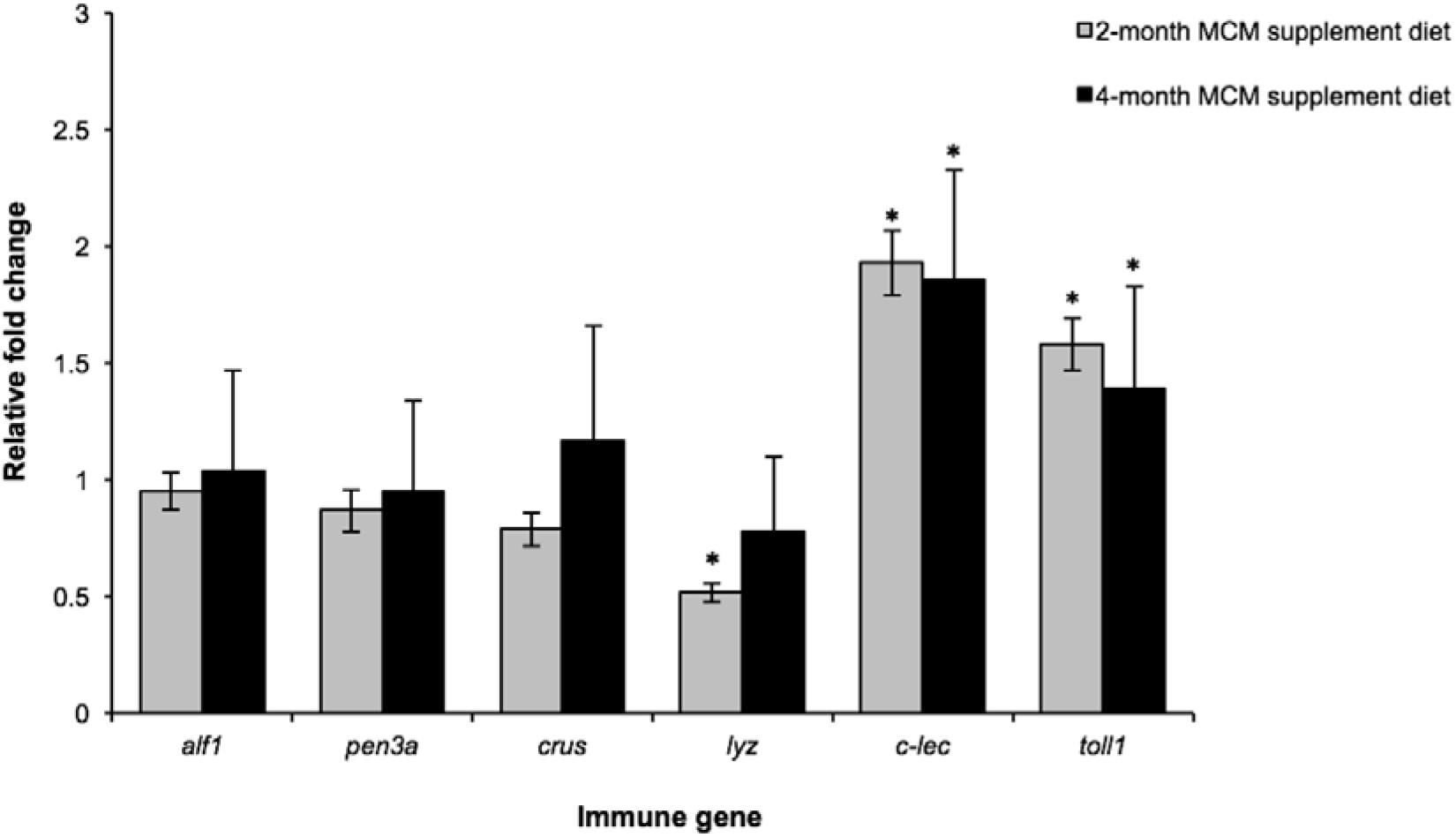
Relative gene expression analysis of immune-related genes in intestines of Pacific white shrimp after 2-month and 4-month feeding period with the mannanase-hydrolyzed coprameal-supplemented diet group (MCM). The fold changes of transcript of antimicrobial peptides *(alf1, pen3a* and *crus)*, lysozyme (*lyz*), C-type lectin *(c-lectin)*, and Toll-like receptor protein *(Toll1)* were determined by real-time PCR in relative to the control diet. An error bar represents standard deviation, which was calculated from five replicates for each sample. Asterisk indicates significant difference in fold-changes between groups fed with MCM and non-MCM containing diets (*p* < 0.05).

Here, we report that the feed supplement containing mannanase-hydrolyzed coprameal (MCM) was able to induce transcript levels of pathogen recognition proteins in the Pacific white shrimp. The significant increase was observable on Month 2 of the feeding trial and the level remained similar on Month 4. Although future shrimp experiments fed with a range of MCM concentration might be necessary to fully elucidate beneficial effects of MCM. Our results provide a promising evidence for further implementation of MCM as feed supplements or immunostimulants to enhance shrimp health. Additionally, a comparison of shrimp survival rates under pathogen challenge between the MCM supplement shrimp group to non-MCM supplement control will strengthen the potential application of MCM as shrimp immunostimulant.

## Acknowledgements

We thank Dr. Wonnop Visessanguan from National Center for Genetic Engineering and Biotechnology (BIOTEC, Thailand) for advice and insights on evaluation of shrimp feed additive. This project was supported the Thailand Research Fund (RSA5780022).

